# Predicting Tumor Cell Response to Synergistic Drug Combinations Using a Novel Simplified Deep Learning Model

**DOI:** 10.1101/2020.04.10.036491

**Authors:** Heming Zhang, Jiarui Feng, Amanda Zeng, Philip Payne, Fuhai Li

## Abstract

Drug combinations targeting multiple targets/pathways are believed to be able to reduce drug resistance. Computational models are essential for novel drug combination discovery. In this study, we proposed a new simplified deep learning model, **DeepSignalingSynergy**, for drug combination prediction. Compared with existing models that use a large number of chemical-structure and genomics features in densely connected layers, we built the model on a small set of cancer signaling pathways, which can mimic the integration of multi-omics data and drug target/mechanism in a more biological meaningful and explainable manner. The evaluation results of the model using the NCI ALMANAC drug combination screening data indicated the feasibility of drug combination prediction using a small set of signaling pathways. Interestingly, the model analysis suggested the importance of heterogeneity of the 46 signaling pathways, which indicates that some new signaling pathways should be targeted to discover novel synergistic drug combinations.

## 1. Introduction

Acquired and innate drug resistance is one major challenge in cancer therapy, due to the complex signaling pathways of cancer. Drug combinations targeting multiple targets or multiple signaling pathways are believed to be one possibility to reduce drug resistance. Many studies have identified to identify potentially effective and synergistic drug combinations for cancer treatment in experimental laboratories. For example, RAS and ERK inhibitors were recently reported to be synergistic with autophagy inhibitors in RAS-driven cancers^1,2^. The mechanism of synergy is that the inhibition of RAS signaling causes the activation of autophagy signaling, which prevents cancer cell death^1,2^. In BRAF inhibitor resistant Melanoma, vemurafenib (BRAF inhibitor) + tretinoin (retinoic acid receptor agonist) were found to be effective and synergistic in cell assays and mouse models. However, there are a few effective drug combinations for clinical use in cancer therapy. Novel and effective drug combinations are needed for personalized treatment to reduce the drug resistance.

Many cancer cell lines and mouse models are available to experimentally screen drugs and drug combinations. However, the experimental screening approaches are limited, considering the numerous possible combinations of thousands of FDA approved drugs and thousands of investigational agents. For example, there are currently about 4 available datasets of experimental screening drug combinations: 1) NCI-ALMANAC Drug Combination Data Set^3^ (∼5,232 combinations from ∼100 drugs on NCI60 cell-lines); 2) the Astraeneca-Sanger Drug Combination Prediction DREAM Challenge Data Set^4^ (900 combinations from 118 compounds on 85 cancer cell lines); 3) the Yale-Stern Melanoma DataSet^5^ (∼7000 combinations from 145 drugs/compounds on 19 melanoma cancer cell lines); and 4) the Merck-2016 DataSet^6^ (583 combinations from 38 drugs/compounds on 39 cancer cell lines). These datasets provide valuable basis to build machine-learning and deep learning-based models.

Computational models that integrate diverse pharmacogenomics datasets with multi-omics data of cancer patients to prioritize drug combinations are essential for novel drug combination discovery. The combination of computational and experimental models can facilitate drug combination discovery in a fast manner. Though a set of prediction models have been reported for drug combination prediction, it remains an open problem. For example, the network-based and connectivity map^7,8^ based drug combination models^9,10^ developed in synergy based on the gene-gene interaction network have been proposed. In addition, a semi-supervised learning model integrating diverse pharmacogenomics datasets was proposed to predict drug combination^11^. Network message propagation-based models developed using drug-target interactions and multi-omics data were also proposed to predict combinations^12,13^. Deep learning models have also been proposed for drug combination prediction. For example, A deep belief network (DBN) model, DeepSynergy, that integrates a large number of chemical structure and genomics features on the Merck-2016 DataSet^6^ was recently proposed^14^ as a prediction drug combination method. The other deep learning model, AuDNNsynergy^15^ (Deep Neural Network Synergy model with Autoencoders), integrates the multi-omics data of over 10,000 cancer genome atlas (TCGA) cancer samples. One limitation of the existing deep learning models of drug combination prediction is the use of a large number of chemical and omics features (>10 thousand features) and fully connected dense layers (a huge number of parameters in the model to be trained) relative to the small number (30∼100 drugs on 30∼60 cancer cell lines) of drug combination synergy scores experimentally obtained. Though the model with a large number of parameters can fit/predict the data, the model parameters cannot be well trained, and cannot not be explained.

To reduce the complicity of the deep learning model and make the models more explainable, in this study, we propose a novel simplified deep learning model, ***DeepSignalingSynergy***, for drug combination prediction. Compared with existing models that make use of a large number chemical and genomics features, we built the model on a small set of cancer signaling pathways, with the aim of investigating the importance of individual signaling pathways. Moreover, the model can mimic the integration of multi-omics data and drug target/mechanism in a relatively more biological meaningful and understandable manner. The results from evaluating the model on the NCI ALMANAC drug combination screening data indicated the feasibility of using a small set of signaling pathways and showed the importance of signaling pathways that affect the drug combination response.

## 2. Materials and Methodology

### 2.1 Drug combination screening data in NCI ALMANAC dataset

The drug pair data was obtained from the NCI ALMANAC database, which is a resource created in 2017. The NCI Almanac dataset includes a score assigned to each of the drug pairs was assigned a score, termed the NCI “ComboScore”^1^ to indicate the synergy scores of drug combinations. In summary, the synergistic effects of combinations of 104 FDA approved drugs in terms of cancer cell growth inhibition were evaluated on NCI 60 cancer cell lines. The average comboScore of two drugs with different doses on a given cancer cell lines was used to indicate the synergy score of two drugs on the cancer cell line, with a 4-element tuple: <*D*_*A*_, *D*_*B*_, *C*_*C*_, *S*_*ABC*_>.

### 2.2 RNA-seq gene expression and copy number data of NCI-60 Cancer Cell Lines from Cancer cell line encyclopedia (CCLE)

Cancer cell line encyclopedia (CCLE) database^16^ provides the multi-omics data of more than 1000 cancer cell lines, e.g., RNA-seq (gene expression), copy number variation, metabolomics, miRNA, RPPA. The large panel of cancer cell lines with comprehensive genetic characterization provide a data source to investigate the associations between molecular features and cancer phenotypes, including drug responses. For this study, the RNA-sequencing gene expression values, using TPM (transcripts per million), and copy number values of genes of 1019 cancer cell lines were downloaded from the cancer cell line encyclopedia (CCLE) website: https://portals.broadinstitute.org/ccle. The CCLE dataset 45 of NCI-60 cancer cell lines were included, as shown in **Table I**.

**Table I:**
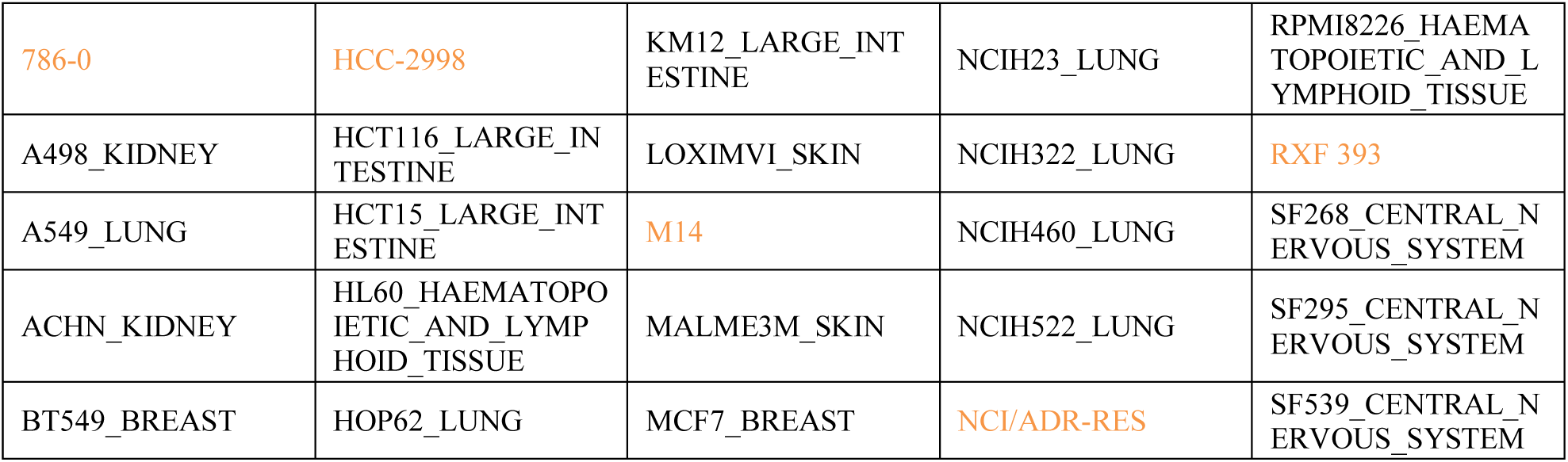
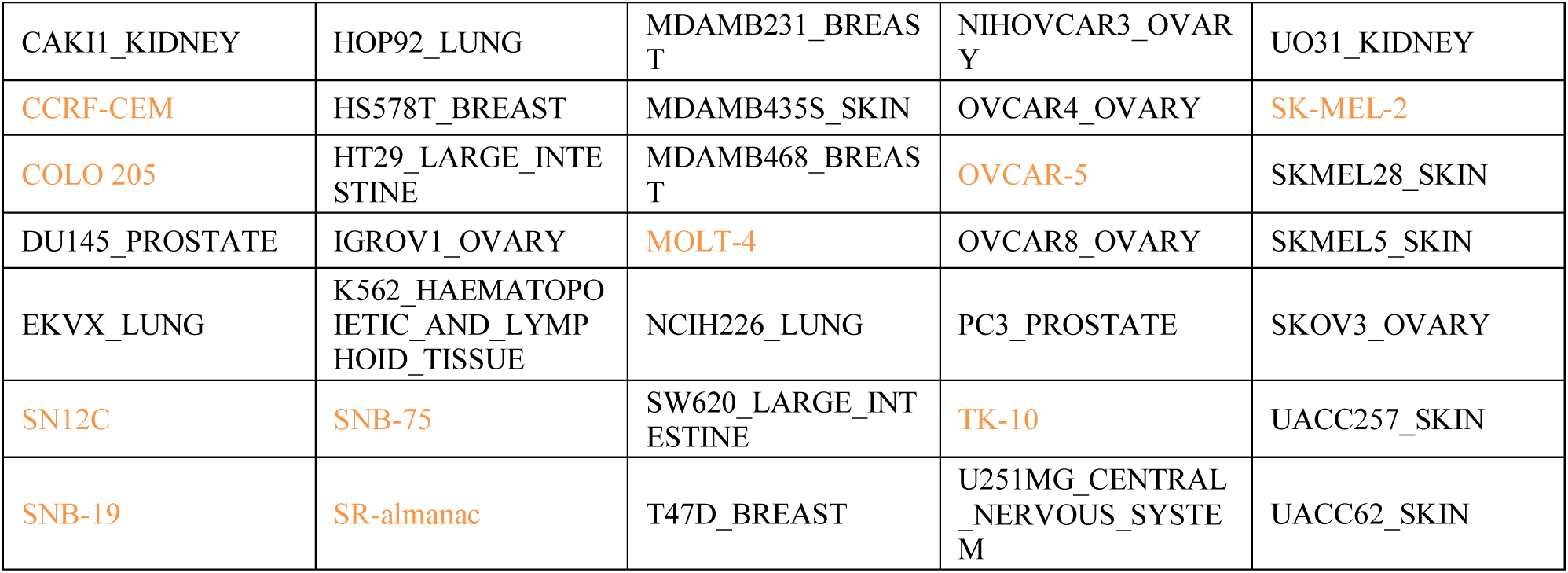
NCI-60 cancer cell lines included in CCLE with RNAseq data. Orange text indicates the cancer cell lines that are not included in CCLE.

### 2.3 KEGG signaling pathways and cellular process

KEGG (Kyoto Encyclopedia of Genes and Genomes)^17^ is a database for the systematic understanding of gene functions. The KEGG signaling pathways provide the knowledge of signaling transduction and cellular processes. There are 303 pathways in KEGG database, and 45 of them are annotated as “signaling pathways”. Many of the signaling pathways are important oncogenic signaling pathways^18^, e.g., EGFR, WNT, Hippo, Notch, PI3K-Akt, RAS, TGFβ, p53. The ‘cell cycle’ cellular process is also included. For simplification, the ‘cell cycle’ is also viewed as one ‘signaling’ pathway. In total, 46 signaling pathways (45 signaling pathways + cell cycle) are selected (see **Table II**). Among these 46 signaling pathways, there are 1648 genes with both gene expression and copy number variation data.

**Table II:** The 46 signaling pathways used in the proposed model.

|  |  |  |  |  |
| --- | --- | --- | --- | --- |
| MAPK | FoxO | TGF-beta | T cell receptor | Adipocytokine |
| ErbB | Sphingolipid | VEGF | B cell receptor | Oxytocin |
| Ras | Phospholipase D | Apelin | Fc epsilon RI | Glucagon |
| Rap1 | p53 | Hippo | TNF | Relaxin |
| Calcium | mTOR | Toll-like receptor | Neurotrophin | AGE-RAGE |
| cGMP-PKG | PI3K-Akt | NOD-like receptor | Insulin | Cell cycle |
| cAMP | AMPK | RIG-I-like receptor | GnRH |  |
| Chemokine | Wnt | C-type lectin receptor | Estrogen |  |
| NF-kappa B | Notch | JAK-STAT | Prolactin |  |
| HIF-1 | Hedgehog | IL-17 | Thyroid hormone |  |

In summary, there are gene expression (TPM) and copy number variation data of 1648 genes in 46 signaling pathways of 45 cancer cell lines, which was used as the input of the deep learning model.

### 2.3 Drug-Target interactions from DrugBank database

DrugBank^19^ is a widely used database to retrieve the information of drugs, such as drug name, chemo-structure, drug mechanism as well as comprehensive drug target information. There are more than 13,000 drug entries in the latest release of DrugBank (version 5.1.5, released 2020-01-03). Among these entries, 2,630 are FDA approved small molecule drugs, and about 6,355 are investigational agents (not approved yet). In total, there are 15263 drug-target interactions between 5435 drugs/investigational agents and 2775 targets. Among the drugs in NCI ALMANAC, 67 drugs are included in DrugBank with targets; and 21 (see **Table III**) drugs with targets on the 1648 signaling pathways were kept as the input of the model.

**Table III:** The 21 drugs used in the proposed model.

|  |  |  |  |
| --- | --- | --- | --- |
| Celecoxib | Gefitinib | Quinacrine hydrochloride | Tretinoin |
| Cladribine | Imatinib mesylate | Romidepsin | Vinblastine sulfate (hydrate) |
| Dasatinib | Lenalidomide | Sirolimus | Vorinostat |
| Docetaxel | Mitotane | Sorafenib tosylate | Thalidomide |
| Everolimus | Nilotinib | Tamoxifen citrate | Paclitaxel |
| Fulvestrant |  |  |  |

### 2.4 Architecture of DeepSignalingSynergy

**Fig. 1** shows the schematic architecture of the proposed *DeepSignalingSynergy* model. In the ‘input layer’, there are 4 input features, i.e., gene expression (RNA-seq TPM values), copy number, is_target_of_*D*_*A*_ (0: this gene is not a target of Drug_A; 1: this gene is a target of Drug_A), and is_target_of_*D*_*B*_, for each of 1648 genes on cancer cell line *C*_*C*_. For the connections between the ‘gene’ and ‘pathway’ layers, the 1648 genes are connected the 46 signaling pathways, only if a gene is included in a signaling pathway (not dense connections). The output of the ‘46 signaling pathway’ is used as the input of the deep belief network (DBN) (densely connected). The ‘output’ layer is the synergy score of a drug combination < *D*_*A*_, *D*_*B*_> on cancer cell line *C*_*C*_. The mean square error (MSE) is used as the loss function. For the DBN, there are 3 hidden layers: first hidden layer has 256 nodes with the relu activation function; the second hidden layer has 128 nodes with the relu activation function; the third hidden layer has 32 nodes with the relu activation function. The linear activation function is used in the output layer.

**Figure 1:**
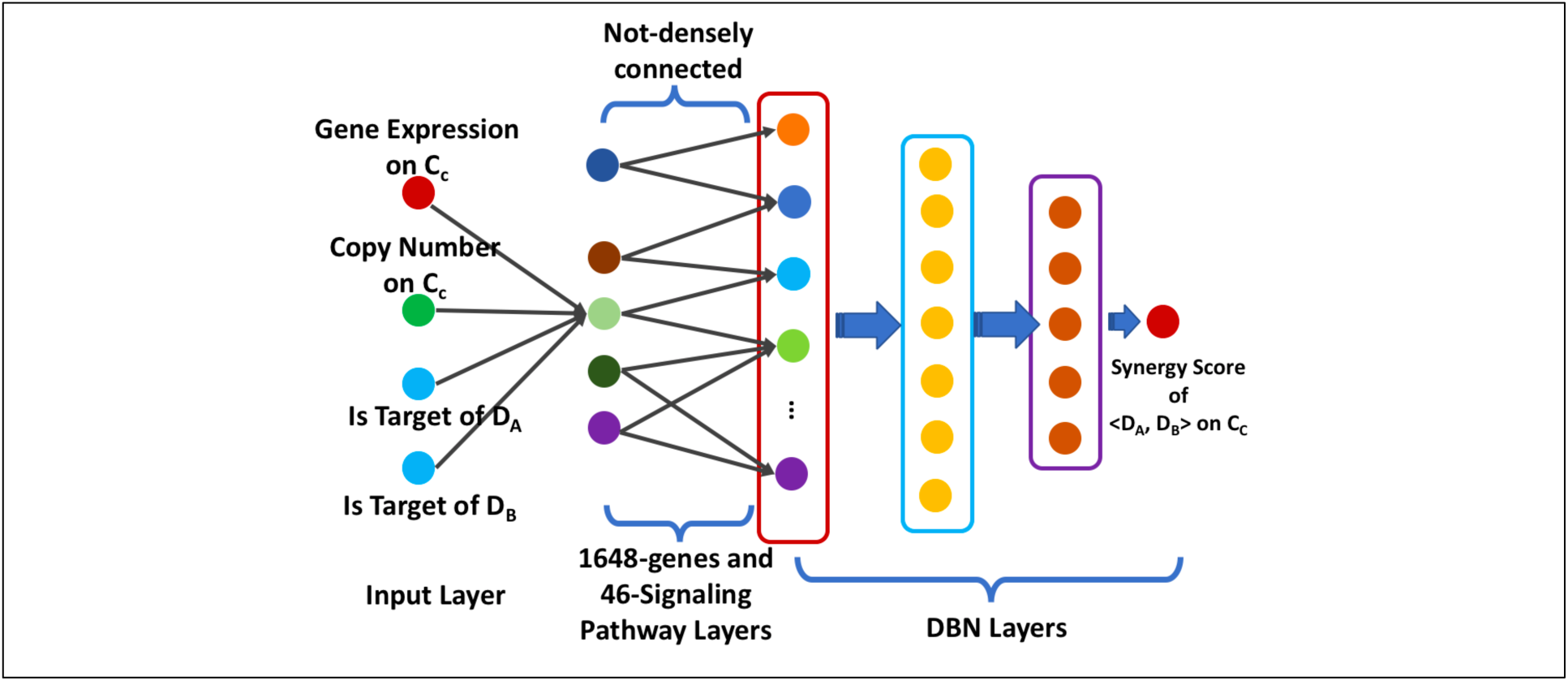
Schematic architecture of the proposed *DeepSignalingSynergy* model.

## 3. Results

### 3.1 Evaluation of drug combination prediction of DeepSignalingSynergy

There are about 5658 synergy scores of 21 drugs on 45 cancer cell lines, i.e., <*D*_*A*_, *D*_*B*_, *C*_*C*_, *S*_*ABC*_>. To evaluate the performance of the DeepSignalingSynergy model, we randomly divided the dataset into a training dataset (80%) and a test dataset (20%) 3 times. The model is trained with 30 epochs. The Pearson correlation was used as the metric for the model performance evaluation. **Fig. 2** shows the evaluation results on 3 randomly selected training and test datasets. The proposed model has the average Pearson correlation coefficients, 0.79 and 0.67 on the 3 training datasets and test datasets respectively (see **Table IV**). This result indicated that the performance of the proposed model, using a small set of signaling pathways, is relatively low but potentially comparable with other existing deep learning models using a large number of chemical-structure and genomics features reported^15^, like AuDNNsynergy^15^ and DeepSynergy^14^ (which have the Pearson correlation coefficients of 0.74 and 0.73 respectively).

**Table IV:** MSEs and Pearson correlation coefficients on the 3 randomly selected training and test datasets.

| Data set | Epoch number | MSE | Pearson Correlation |
| --- | --- | --- | --- |
| Training-1 | 30 | 35.6 | 0.769 |
| Test-1 | 30 | 41.3 | 0.673 |
| Training-2 | 30 | 32.3 | 0.801 |
| Test-2 | 30 | 42.8 | 0.674 |
| Training-3 | 30 | 32.0 | 0.804 |
| Test-3 | 30 | 41.9 | 0.666 |

**Figure 2:**
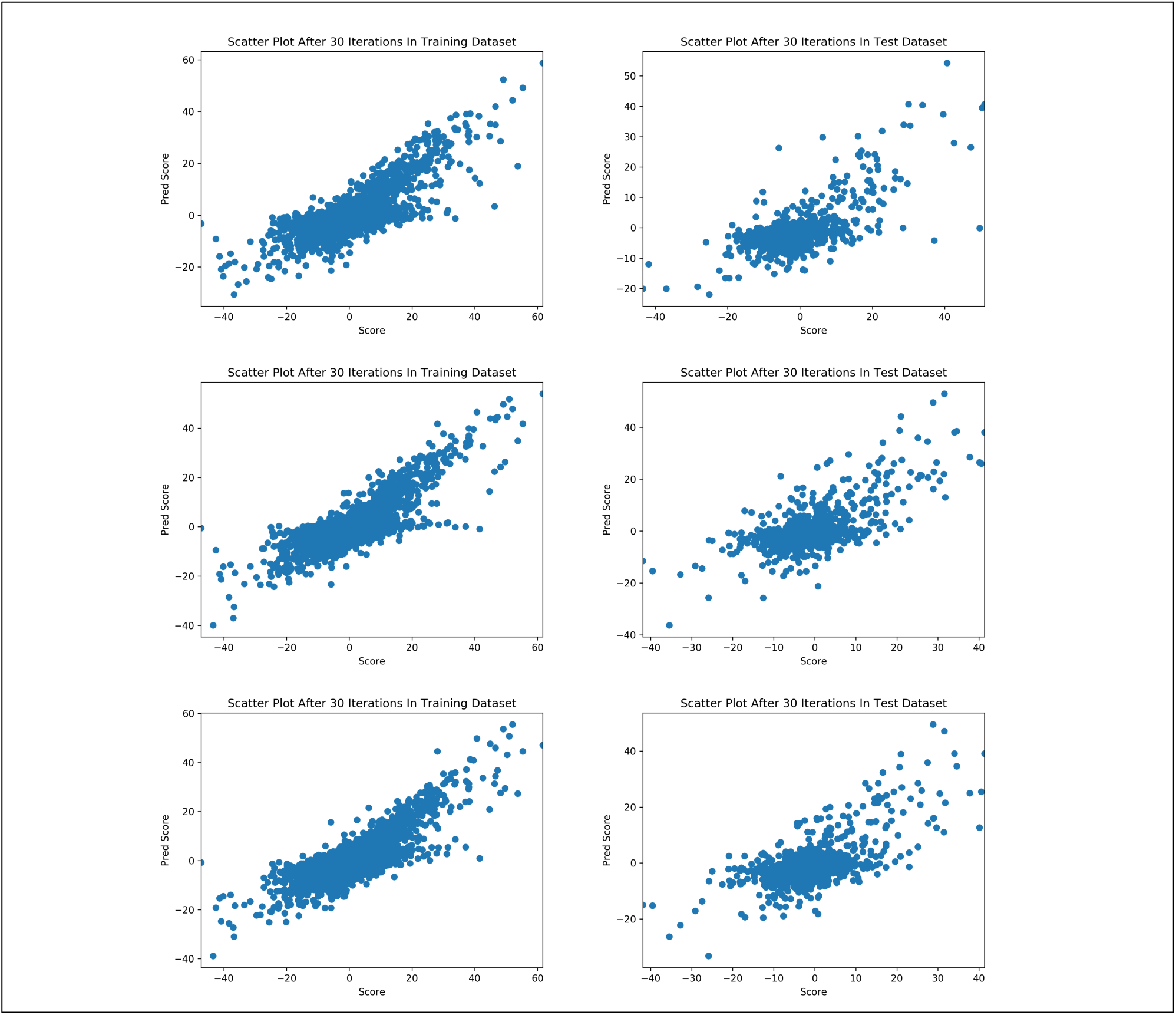
Scatter plot of the predicted and experimental synergy scores at 40 epoches of 3 randomly selected training (80%) and test (20%) datasets respectively.

### 3.2 Importance of signaling pathways analysis for understanding potential mechanism of synergy

To investigate the potential mechanism of synergy in terms of the contributions of individual signaling pathways to the prediction of synergy of drug combination, we employed the Layer-Wise Relevance Propagation (LRP) approach implemented in the “iNNvestigate” package^20^, which can be used to visualize the importance of individual inputs at different layers. **Fig. 3 and Fig. 4** show the density distribution maps of importance scores of 46 signaling pathways over the 3 randomly selected test datasets. The results indicated that the importance of the individual signaling pathways are relatively stable in the 3 randomly selected test datasets. Though the importance scores are positive or negative in different test datasets, the rough range and values of absolute importance scores are consistent. First, interestingly, some of the 46 signaling pathways, e.g., the MAPK, TGF-β, cell cycle, AMPK, RAS, Jak-Stat, HIF-1α signaling pathways have much more importance than other oncogenic signaling pathways. Second, the Apelin, Adipocytokine, Fc epsilon RI, Neurotrophin, and IL-17 signaling pathways surprisingly contribute to the drug combination response prediction. Third, some signaling pathways showed the similar interesting distributions, e.g., the MAPK and RAS signaling pathways, the FoxO and cAMP signaling pathways, as well as Apelin and Neurotrophin signaling pathways. Fourth, other oncogenic signaling pathways, like the mTOR, ER, Hippo, Rap1 signaling pathways, can only contribute to the drug combination synergy prediction moderately. Though results are interesting, further investigations are needed to understand and explain the roles of individual signaling pathways and their associations with drug combination response.

**Figure 3:**
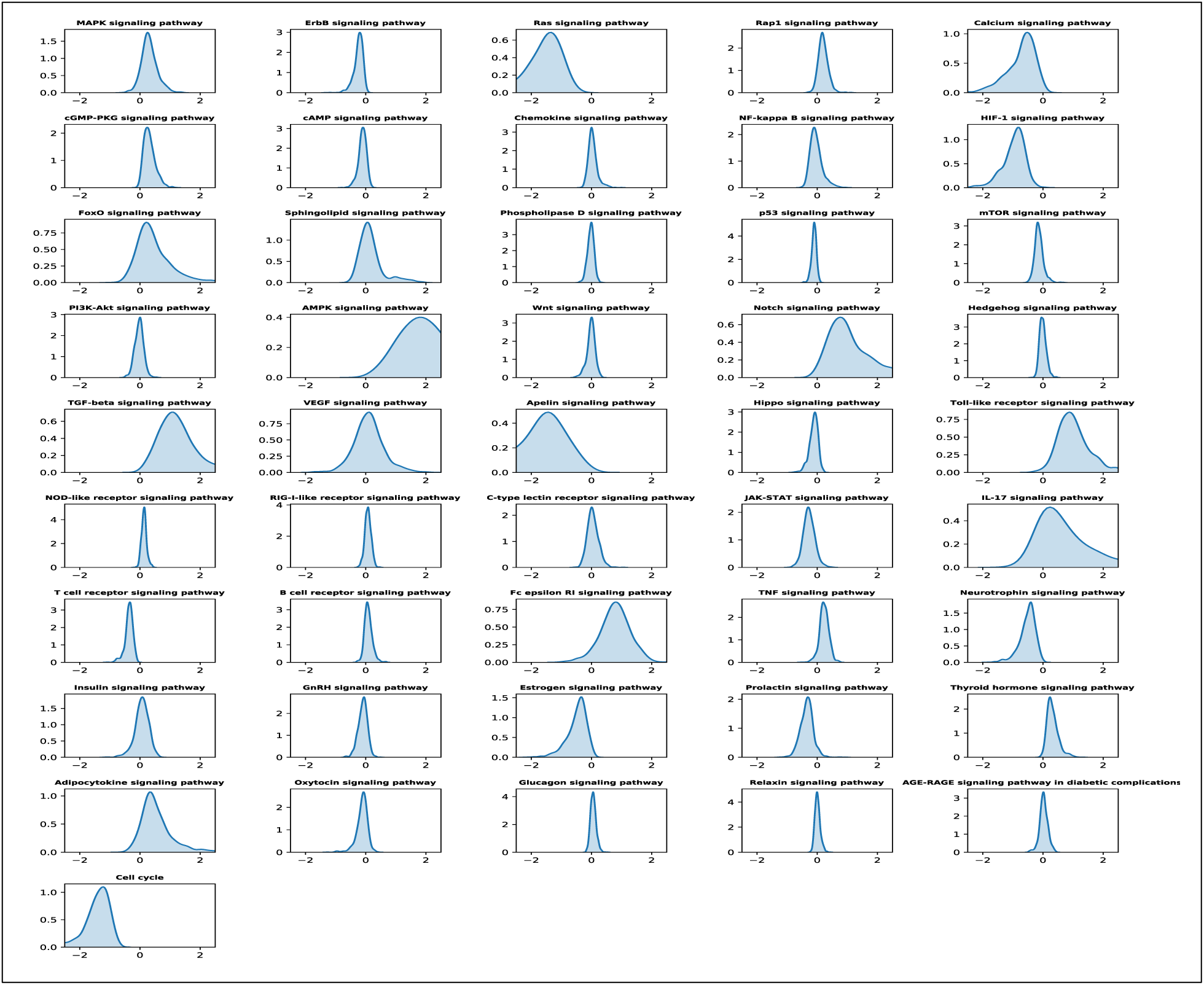
Distribution of importance of 46 signaling pathways on the first test dataset.

**Figure 4:**
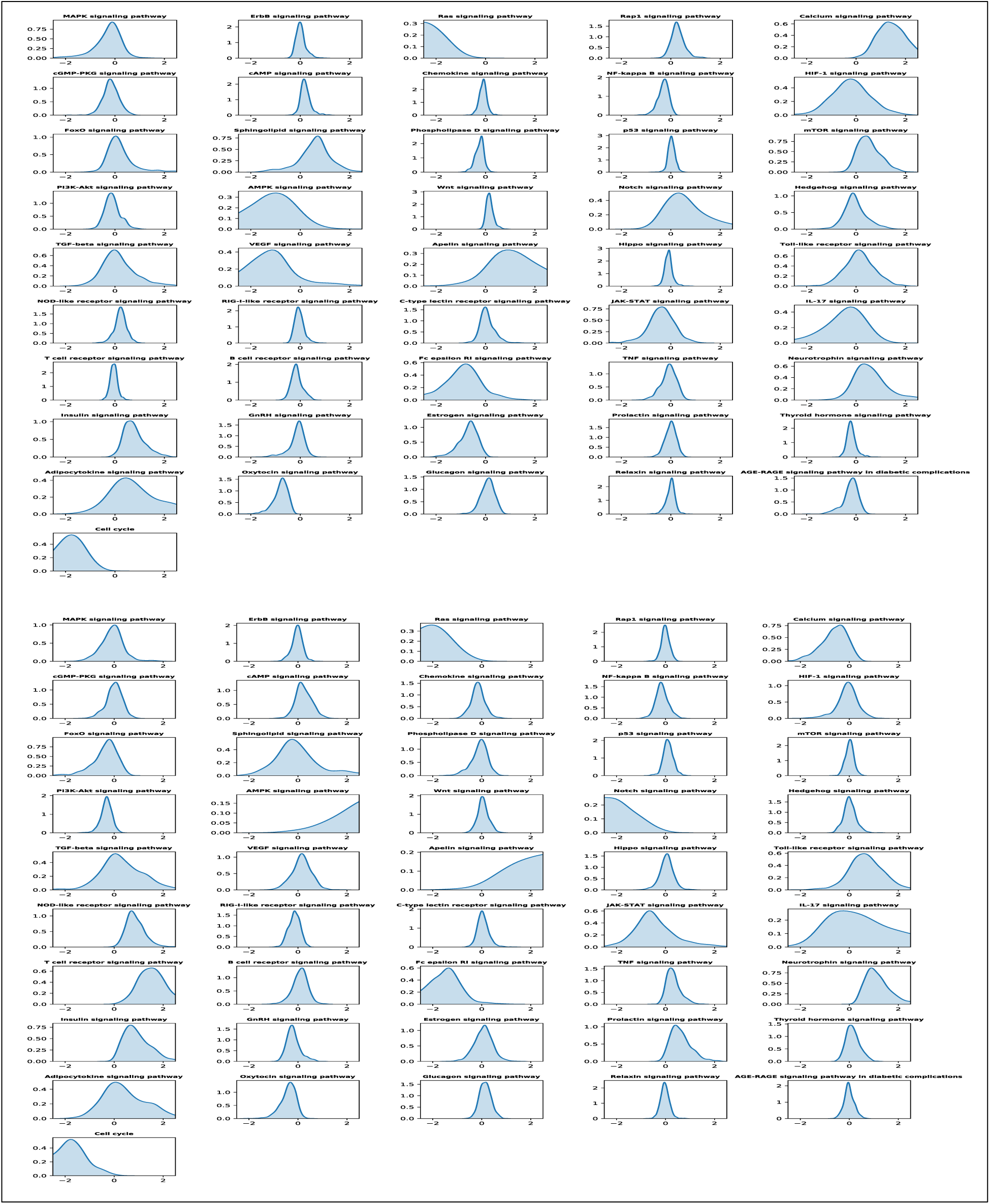
Distributions of importance of 46 signaling pathways on the second (top-panel) and third (bottom-panel) test datasets respectively. The results indicated that the importance of the individual signaling pathways are relatively stable in the 3 randomly selected test datasets. Though the importance scores are positive or negative in different test datasets, the rough range and values of absolute importance scores are consistent.

### 3.3 Importance of individual genes

We conducted the similar analysis to investigate the importance of individual genes. **Fig. 5** shows the top 50 genes with the largest absolute importance scores of 1648 genes on the 3 randomly selected test datasets respectively. The selected top 50 genes (out of 1648 genes, which is much more than 46 signaling pathways) are not so consistent. The common genes selected in all the 3 test datasets are: ‘PRKCG’, ‘KIT’, ‘RRM2B’, ‘FLT3’, ‘PDGFRA’, ‘PDPK1’, ‘JUN’, ‘NTRK1’, ‘BCL2’’, which indicate the potential synergy among these targets. It can be possible to understand the mechanism of synergy further by investigating the importance scores of individual genes and pathways for a specific synergy drug combination on a specific cancer cell line. However, it is still challenging to associate the importance scores to specific synergy mechanism of drug combinations.

**Figure 5:**
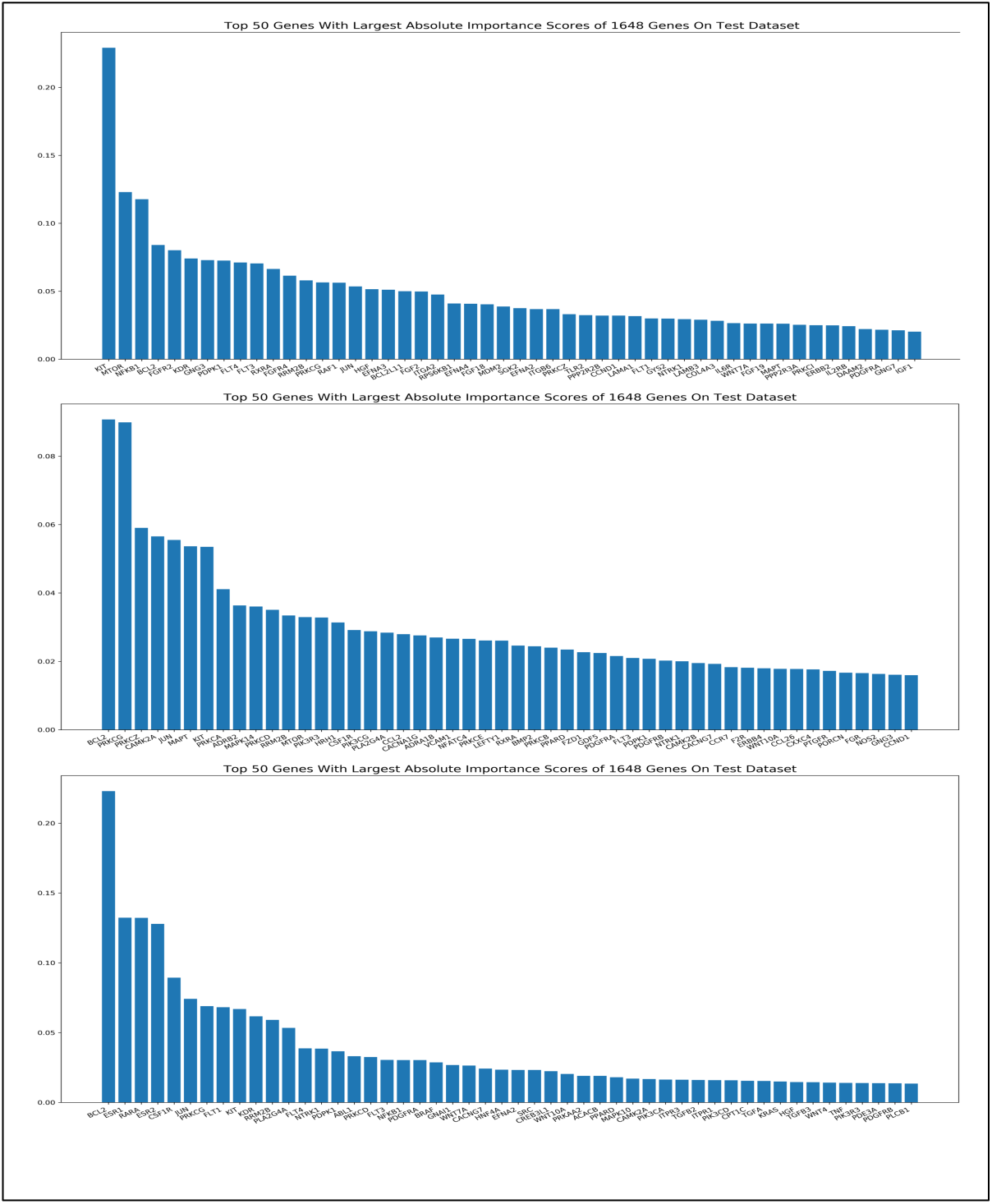
Importance of individual genes.

## 4. Discussion and conclusion

Synergistic drug combinations are important factors in reducing drug resistance in cancer therapy. Computational models that can integrate multi-omics data of cancer patients with pharmacogenomics data of drugs and investigational agents are needed to predict potential synergistic drug combinations (to narrow down the search space of drug combination). The combination of computational and experimental models can facilitate the discovery of synergistic drug combinations in a fast manner.

Deep learning models have been widely used and outperform the traditional machine learning models in image analysis, natural language processing, healthcare data analysis, and drug combination prediction. However, it is a challenging to make the model explainable, especially the models with a large number of features and parameters. In the existing deep learning models of drug combination prediction, a large number of chemical-structure and genomics features are used via the densely connected layers, which requires the training of a large number of parameters. However, only small sets of drug combination experimental validation results that can be used as training labels are available. Thus, it is hard to train the large number of parameters well, and it is also hard to explain the model to investigate the potential mechanism of drug combination synergy.

In this study, we propose to reduce the number of parameters in deep learning models by using a simplified deep learning model built based on a set of biological meaningful signaling pathways. In the model, we can integrate multi-omics data of individual genes and drug-target information, and link the genes to 46 pathways in a sparse manner with a much fewer number of parameters (compared to densely connected layers). The evaluation results showed that the proposed simplified model can achieve good prediction results in terms of Pearson correlation coefficient between the predicted and experimental synergy scores. Moreover, the explainable analysis of the deep learning model identified some interesting results in terms of the importance of individual signaling pathways that contribute to the drug combination synergy. Further analyses are needed to investigate the unclear mechanisms of synergy using these signaling pathways.

This is our first expletory study to investigate and prediction drug combination synergy with a simplified deep learning model with increased possibility of model explanation. There are some limitations of the proposed model that need to be further addressed. First, STITCH^21^ database can provide much more drug-target interactions, in addition to drug-target interactions obtained from DrugBank. With more drug-target interactions, more drugs can be included to the model, and the prediction accuracy could be better. Second, in addition to the 46 signaling pathways, other KEGG pathways, like metabolism pathways, will be further evaluated. Third, Gene oncology^22^ (GO) terms provide alternative biological meaningful biological processes (BP) (gene sets), which can cover many more genes (drug targets) and can be used for drug combination prediction. Third, other omics data, like protein, methylation, genetic mutation can be integrated conveniently to the model in addition to the copy number, gene expression data. We will investigate these possible directions in the future work. Moreover, we will develop novel approaches to uncover the explainable mechanisms of synergy of drug combinations, e.g., the synergy mechanism of RAS/ERK inhibitors and Autophagy inhibitors recently reported^1,2^, which can provide clues to discover novel synergistic drug combinations to reduce drug resistance in cancer therapy.

